# Social boldness correlates with brain gene expression in male green anoles

**DOI:** 10.1101/2021.01.15.426859

**Authors:** David Kabelik, Allison R. Julien, Dave Ramirez, Lauren A. O’Connell

**Author notes:** Please send correspondence to: David Kabelik, Department of Biology, Rhodes College, 2000 N Parkway, Memphis, TN 38112, USA.

## Abstract

Within populations, some individuals tend to exhibit a bold or shy social behavior phenotype relative to the mean. The neural underpinnings of these differing phenotypes – also described as syndromes, personalities, and coping styles – is an area of ongoing investigation. Although a social decision-making network has been described across vertebrate taxa, most studies examining activity within this network do so in relation to exhibited differences in behavioral expression. Our study instead focuses on constitutive gene expression in bold and shy individuals by isolating baseline gene expression profiles that influence social boldness predisposition, rather than those reflecting the results of social interaction and behavioral execution. We performed this study on male green anole lizards (*Anolis carolinensis*), an established model organism for behavioral research, which provides a crucial comparison group to investigations of birds and mammals. After identifying subjects as bold or shy through repeated reproductive and agonistic behavior testing, we used RNA sequencing to compare gene expression profiles between these groups within various forebrain, midbrain, and hindbrain regions. The ventromedial hypothalamus had the largest group differences in gene expression, with bold males having increased expression of neuroendocrine and neurotransmitter receptor and calcium channel genes compared to shy males. Conversely, shy males express more integrin alpha-10 in the majority of examined regions. There were no significant group differences in physiology or hormone levels. Our results highlight the ventromedial hypothalamus as an important center of behavioral differences across individuals and provide novel candidates for investigations into the regulation of individual variation in social behavior phenotype.

## Introduction

Individuals vary widely in their social boldness. Some individuals perform many high intensity behaviors within moments of participating in a novel social encounter, while others hesitantly engage in a few low-intensity interactions. Often, such social boldness is consistent across different social environments, and correlates with other behavioral traits like active versus passive stress coping (Coppens et al., 2010; Koolhaas et al., 2010; Stamps and Groothuis, 2010). Although a continuum of such behavioral propensity usually exists within a population, we can categorize individuals at each end of such a continuum as either behaviorally ‘bold’ or ‘shy’, with bold individuals exhibiting lower latency, higher frequency, and higher intensity of exhibited social behaviors across contexts than shy individuals. Such behavioral phenotypes, also referred to as behavioral syndromes, personalities, or coping styles (Koolhaas et al., 2010; Réale et al., 2010; Sih et al., 2004), often manifest as correlated suites of behavioral outputs, presumably due in part to regulation by shared neural underpinnings. The neural substrates that lead an individual toward exhibiting a bold or shy phenotype likely rely on brain regions involved in social decision-making (Newman, 1999; O’Connell and Hofmann, 2012, 2011) and neuroendocrine mediators of these circuits (Baugh et al., 2012; Félix et al., 2020; Ketterson and Nolan Val, 1999). Although numerous neural systems have been associated with social behavioral output, the specific proximate variables that determine stable bold-shy behavioral phenotypes remain unknown. Understanding how the brain regulates social behavior propensity is both fundamentally intriguing and also necessary to address various disorders involving social anxiety, depression, and aggression.

Among vertebrates, the lack of understanding of neuroendocrine regulators of behavioral phenotypes is especially true among non-avian reptiles, as they are the least studied vertebrate taxonomic group (Kabelik and Hofmann, 2018), despite serving as an important evolutionary comparison group, especially for amniotic vertebrates. A social decision-making network has been described in reptiles (Kabelik et al., 2018), and various neuroendocrine variables have been related to the expression of social behaviors in lizards (Dunham and Wilczynski, 2014; Hartline et al., 2017; Kabelik et al., 2013, 2008b; Kabelik and Crews, 2017; Kabelik and Magruder, 2014; Korzan et al., 2001; Korzan and Summers, 2004; Larson and Summers, 2001; Smith and Kabelik, 2017; Watt et al., 2007; Woolley et al., 2004a, 2004b, 2001). However, many potential neural, endocrine, and direct genetic regulators of social boldness remain unexamined. In this study, we compare gene expression from various brain regions of male green anoles (*Anolis carolinensis*) that exhibit either stable bold or shy phenotypes in order to identify potential regulatory variables. Green anoles are a longstanding model for social behavior investigation (Lovern et al., 2004), and they have recently become a model for comparative genomic investigation (Alföldi et al., 2011), making them an ideal subject species for the present study. We focus here on male green anoles because they exhibit high levels of both reproductive and aggressive behaviors, and our aim was to differentiate individuals based on boldness within both contexts.

Many studies of social boldness examine gene expression resulting from the performance of specific social behaviors (e.g., Mukai et al., 2009; Wong et al., 2012; Zayed & Robinson, 2012), or by adoption of a dominant or submissive status within a social hierarchy (e.g., Eastman et al., 2020; Renn et al., 2008). Here we instead examine differences in baseline neural gene expression among subjects that have been extensively screened within different social contexts and assigned to a bold or shy phenotype category. This design eliminates gene expression differences associated with expressed behavioral output and instead places focus on the neural state differences that may be associated with consistent bold or shy behavioral outputs outside of behavioral encounters with conspecifics. Additionally, the examined males are housed individually and thus hold identical home ‘territories’, eliminating social status-related gene transcription. We selected five bold and five shy individuals for our transcriptome analysis and compared gene expression profiles between these experimental groups across several brain regions.

As bold and shy behavioral traits have been associated with brain regions governing social behavior in other taxa (Delclos et al., 2020; Geng and Peterson, 2019; Koolhaas et al., 2010), we examined gene expression in four forebrain, one midbrain, and one hindbrain region of the social behavior network. The examined brain regions include nodes of the social decision-making network, such as the preoptic area, lateral septum, and ventromedial hypothalamus of the forebrain, as well as the ventral tegmental area of the midbrain. Additional regions that we sampled included the medial and dorsomedial cortex, which are considered at least partly homologous to the mammalian hippocampal complex (Desfilis et al., 2018; Striedter, 2016; Tosches et al., 2018), and the dorsal ventricular ridge, an area partly homologous and partly analogous to the mammalian neocortex (Briscoe et al., 2018; Briscoe and Ragsdale, 2018; Colquitt et al., 2021; Northcutt, 1981) and amygdaloid nuclei (Lanuza, 1998; Martínez-García et al., 2002). Finally, a rostral hindbrain region was also examined, as the hindbrain has also been linked to the regulation of social behaviors (Thompson et al., 2008; Walton et al., 2010). We tested the hypothesis that bold and shy male green anoles would differ in sex steroid hormone levels and gene expression, with an emphasis on hypothalamic regions.

## Materials and Methods

### Subjects

Fifty-seven focal male green anoles (*Anolis carolinensis*) were obtained from a commercial supplier. These males were housed singly within terraria (30.5 cm H x 26 cm W x 51 cm L) and kept in breeding season conditions: long-day (14 light:10 dark) full-spectrum lighting, 12 hours of supplemental heat provided 5 cm above one end of a wire-mesh terrarium lid by means of a 60-W incandescent light bulb, and thrice-weekly feeding with crickets. Additional males and females from our housing colony were used in social interactions. All procedures involving live animals were conducted according to federal regulations and approved by the Institutional Animal Care and Use Committee at Rhodes College.

### Social behavior boldness assessment

Behavioral trials were carried out from May 14 to July 3, 2013. Focal males were each assessed three times with different conspecifics for social boldness within each of three social encounter scenarios – reproductive encounter, agonistic encounter as a resident, and agonistic encounter as an intruder. Thus, each focal male’s behavior was scored in nine separate 10-min behavioral encounters, and a maximum of one social encounter per focal male was run per day. The *reproductive behavior* scenario involved two conspecific adult females simultaneously placed into the focal male’s terrarium. Two females were used to maximize the probability of eliciting reproductive behaviors from the focal male. We recorded the frequency (sum of behaviors per 10-min session) and latency to first performance (minute of first occurrence of any listed behavior) of the following behaviors: head bob bout, push-up bout, dewlap extension bout, dewlap extension bout with push up, chase, and copulate. Focal males that failed to display any behaviors were assigned the maximum latency score of 10 min. The maximum intensity of behavioral display was also scored from 0-3 based on the highest achieved category: no display, display only, chase, and copulate. The *agonistic encounter as a resident* scenario involved a size-matched (within 3 mm snout-vent length) adult conspecific male intruder being placed within the focal male’s terrarium. Behaviors were scored as in the reproductive encounter, except that biting of the stimulus male replaced copulation as the highest intensity behavior. The *agonistic encounter as an intruder* scenario involved the focal male being taken out of his terrarium and placed into the terrarium (home territory) of a size-matched adult conspecific male. Behavioral scoring was the same as in the previous agonistic scenario. Stimulus animals were also only used once per day, and no behavioral trials involved the repeated pairing of the same subjects.

### Bold-shy categorization

We conducted principal components analysis (PCA) using SPSS Statistics 22 (IBM) to reduce the average behavioral latency, frequency, and intensity scores from each of the three social behavior interaction scenarios into a single value. For example, in male-female trials, the behavioral latency, frequency, and intensity scores for each male were averaged across the three trials in which he took part, in order to generate an “average reproductive latency”, “average reproductive frequency”, and “average reproductive intensity” score. These average scores were then included in the PCA. In each scenario, the resulting analysis generated a single PCA axis with an eigenvalue > 1, and in each case, this axis was highly positively correlated with average frequency and intensity scores, and negatively with average latency scores (r>±0.73, p<0.001 for each). This PCA axis 1 explained 65% of the behavioral variation in the reproductive boldness trial, 79% of the variation in the agonistic trial as resident, and 80% of the variation in the agonistic trial as intruder (see the supplementary materials for scree plots and loading tables). We used these PCA axes to correlate boldness across behavioral scenarios. We also took an average of these three PCA axes to use in selecting bold and shy individuals for the RNAseq portion of this study. Because the average PCA axis 1 score differed across the three behavioral testing blocks (F(2,54)=4.23, p=0.02), we ranked focal males based on this average PCA principal component axis 1 score within each behavioral block. We then chose the highest and lowest scoring focal male within each block, as well as the next highest and next lowest scoring focal male in two of the three blocks. This resulted in selection of the five most socially ‘bold’ and five most ‘shy’ males out of the 57 focal males screened for behavioral consistency. Scatterplots of PCA scores were made with ggplot2 (version 3.3.0) in RStudio (version 1.3.1056) running R (version 3.5.2).

### Tissue harvesting and brain tissue punching

Prior to handling for blood and brain harvesting, focal subjects were left undisturbed in their home terraria for 1-3 days following their last behavioral trial. We euthanized focal males by cutting through the spinal column and immediately collected trunk blood for hormone analyses (average collection time from first handling was 162 ± 3.2 s). The blood was kept at 4°C until centrifugation. The brain was then rapidly dissected, placed within a microcentrifuge tube filled with Tissue Tek (Sakura) cutting medium, and frozen under dry ice (average time from first handling to freezing of brain was 544 ± 8.4 s). The body (minus the head and some blood) was then weighed, after which the testes were dissected from the body and also weighed. Brains were sectioned at 100 µm on a Microm HM 520 cryostat (Thermo Scientific). The sections were laid onto glass microscope slides resting on a metal block within the cryostat at −19°C. Tissue punches of selected areas were obtained using a Stoelting brain punch set with the aid of a dissecting microscope (Olympus SZX7) mounted above the cryostat. Brain punches were placed into Trizol (Invitrogen) and frozen at −80°C. Either 1 mm or 1.25 mm tissue punches were used to obtain tissue from selected brain regions (see Supplementary Figure 1 for brain punch locations). These brain punch locations were as follows: POA-LS, a region including the preoptic area, anterior hypothalamus, paraventricular nucleus of the hypothalamus, and septal nuclei; HIP, a region of the medial and dorsomedial cortices, which are at least partly homologous to the mammalian hippocampus (Desfilis et al., 2018; Striedter, 2016; Tosches et al., 2018); DVR, including the subcortical pallium (dorsal ventricular ridge, including amygdaloid nuclei) as well as striatum; VMH, the ventromedial hypothalamus; MID, the midbrain tegmentum; HIND, the pons and rostral medulla, though not cerebellum. Brain regions were determined by reference to multiple atlases and publications (Bruce and Braford, 2009; Butler and Hodos, 2005; Greenberg, 1982; Hoops et al., 2018; Jarvis, 2008; Kabelik et al., 2014; Lopez et al., 1992; Naik et al., 1981; O’Connell and Hofmann, 2011; Rosen et al., 2002; ten Donkelaar, 1998).

**Figure 1.**
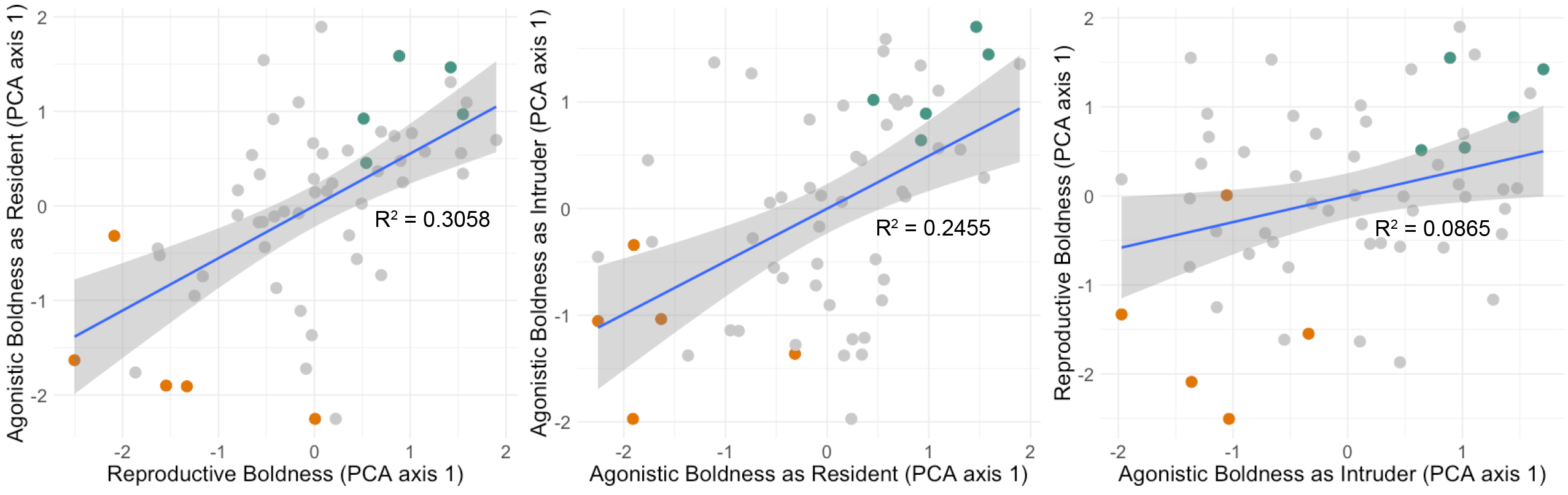
Focal male green anole lizards exhibit stable social boldness phenotypes. (Left) Average boldness in three reproductive encounters correlates strongly with average boldness as the resident male in three separate agonistic encounters. (Middle) Average boldness as the resident male correlates strongly with average boldness as an intruder within three separate agonistic encounters. (Right) Average boldness as the intruder male in three separate agonistic encounters correlates weakly with average reproductive boldness to three pairs of females. Social boldness is represented by PCA axis 1 values, which are positively correlated with average behavioral frequency and intensity, and negatively correlated with average latency to display, within each behavioral encounter scenario. Trials were carried out in three blocks. One to two focal males with the highest combined PCA values per block (‘bold’, shown in blue-green), and one to two focal males with the lowest combined PCA values per block (‘shy’, shown in orange) were selected for the bold-shy neural RNA sequencing comparison.

### Hormone analyses

Blood samples were centrifuged and plasma (averaging 77 ± 3.3 µl) was frozen at −80°C until hormone analysis. We quantified testosterone (ADI-900-065; sensitivity 5.67 pg/mL), estradiol (ADI-900-008; sensitivity 28.5 pg/mL), progesterone (ADI-900-011; sensitivity 8.57 pg/mL), and cortisol (ADI-900-071; sensitivity 56.72 pg/mL) using enzyme-linked immunosorbent assay (ELISA) kits (Enzo Life Sciences, Farmingdale, NY). The cortisol kit cross-reacts with corticosterone at 28%, representing a general glucocorticoid assay, albeit with lower-than-typical sensitivity. Samples were run across two plates for each hormone and the inter-assay variation across plates and the intra-assay variance for each plate is as follows: testosterone (inter: 5.6%; intra: 5.6% and 6.1%), estradiol (inter: 4.1%; intra: 3.7% and 8.5%), progesterone (inter: 4.7%; intra: 2.3% and 4.7%), and cortisol (inter: 6.4%; intra: 3.8% for both plates). We re-suspended 7 µl of plasma in 203 µL of the appropriate assay buffer and ran each sample in duplicate as per manufacturer’s instructions. Hormone results were generally consistent with previously reported levels in this species (Greenberg and Crews, 1990).

### RNA sequencing and analyses

Brain RNA was extracted using Trizol according to manufacturer’s instructions (Thermo Fisher Scientific). Poly-adenylated RNA was isolated from each sample using the NEXTflex PolyA Beads purification kit (Perkin Elmer). Strand-specific libraries with unique barcodes were prepared using the NEXTFLEX Rapid Directional RNA-Seq kit 2.0 according to manufacturer’s instructions (Perkin Elmer). Libraries were pooled in equal molar amounts and sequenced on an Illumina HiSeq 2500 to obtain roughly 40 million reads per sample.

We first applied quality and adaptor trimming to the raw reads using Trim Galor! (http://www.bioinformatics.babraham.ac.uk/projects/trim_galore/; parameters: trim_galore -- paired --phred33 --length 36 -q 5 --stringency 5 --illumina -e 0.1). Reads were then aligned using kallisto (Bray et al., 2016) with default parameters to the *Anolis carolinensis* cDNA reference transcriptome (Anolis_carolinensis.AnoCar2.0.cdna.all.fa.gz) downloaded from Ensembl (May 2020). Read counts were combined into a single matrix. Differences in gene expression within each brain region were calculated using DESeq2 (Love et al., 2014) within the *in silico* Trinity pipeline (p<0.05, 4-fold change). We corrected p-values for multiple hypothesis testing and considered transcripts with false discovery rate (FDR) correct p-values <0.05 significantly differentially expressed. We performed a gene ontology enrichment analysis for differentially expressed genes in the ventromedial hypothalamus using the PANTHER (version 14; http://pantherdb.org/; Mi et al., 2019). Data visualizations were made in RStudio (version 1.3.1056) running R (version 3.5.2). The PCA analysis for gene expression was performed using the prcomp function in the R base package on normalized gene counts and the PCA was plotted using the fviz_pca_ind function in factoextra (version 1.0.7). We tested for brain region and behavioral group differences in principal components using the aov function in the R base package. Boxplots and bar charts were made with ggplot2 (version 3.3.0), the volcano plots were generated using EnhancedVolcano (version 1.0.1), and the heatmap was generated with heatmap.2 function in gplots (version 3.1.1).

### Statistical analyses

Some behavioral scores and all hormone levels were ln-transformed to meet assumptions of parametric analyses. Data reduction was conducted using PCA, and the comparison of the average resultant score across behavioral testing blocks was conducted using one-way analysis of variance. Correlations among behavioral scores were conducted using Pearson’s r. Behavioral and physical differences between bold and shy focal males were examined via independent-samples t-tests, except for behavioral intensity measure, which was compared using a Mann-Whitney U test. Analysis of gene expression of candidate genes from the literature was performed by obtaining the normalized gene expression for each gene and using a linear mixed model in R (lmer function in the lme4 package, version 1.1–21) to test for differences among behavioral group, brain regions, and their interaction with individual as a random effect to account for repeated sampling of brain regions from the same lizards. Significant differences were then followed up with pairwise comparisons in R using emmeans (version 1.4.6), which adjusts for multiple comparisons.

## Results

### Correlated behavioral traits are stable within individuals

Individual differences in social boldness are relatively stable across different types of social encounters. In **Figure 1**, we present correlations between focal males in the reproductive, agonistic as a resident, and agonistic as an intruder scenario, reflecting latency, frequency, and intensity measures reduced into a single principal component axis for each scenario. We found positive correlations between boldness scores across all behavioral scenarios: reproductive boldness and boldness as the resident in an agonistic trial (r=0.55, N=57, p<0.001), between boldness as the resident in an agonistic trial and boldness as the intruder in an agonistic trial (r=0.50, N=57, p<0.001), and between reproductive boldness and boldness as the intruder in an agonistic trial (r=0.29, N=57, p=0.026).

### Behavioral, but not physiological, traits differ between bold and shy individuals

Relative to shy males, the bold males showed a higher average frequency of reproductive behaviors, agonistic behaviors as a resident, and agonistic behaviors as an intruder (**Table 1**). Similarly, the bold males exhibited lower average latencies to first reproductive behavior, to first agonistic behavior as a resident, and to first agonistic behavior as an intruder. Bold males also exhibited higher average behavioral intensities to females, as resident males in agonistic trials, and as intruder males in agonistic trials. These results confirm that our categorization of subjects as bold or shy effectively discriminated individuals upon all three boldness measures (latency, frequency, and intensity). However, behavioral boldness was not correlated with physical (p ≥ 0.10 for all) or hormonal (p ≥ 0.28 for all) characteristics across all 57 males, nor between bold and shy groupings (**Table 1**). Bold and shy focal males did not differ in snout-vent length, or body-minus-head mass (the body was weighed after blood sampling and brain removal, so as not to delay freezing of brain tissue). Likewise, these groups did not differ in testes mass, or in circulating testosterone, estradiol, progesterone, or glucocorticoid levels.

**Table 1.** Mean and standard error (S.E.) values for behavioral and physical variables of bold and shy male green anoles. No physical but all behavioral variables differ between bold and shy groups. Parametric comparisons state the t statistic (t) and degrees of freedom (df); nonparametric comparisons state the *U* statistic (*U*) and the sample size (N); both state the probability of significance (P) at α=0.05.

| | Shy<br>mean | Shy<br>S.E. | Bold<br>mean | Bold<br>S.E. | $t/U$ | Df/N | P |
| --- | --- | --- | --- | --- | --- | --- | --- |
| **Behavioral Variables:** |  |  |  |  |  |  |  |
| reproductive behavior frequency<br>(#/10 min) | 11.1 | 2.62 | 27.9 | 1.72 | -5.36 | 8 | **0.001** |
| resident agonistic frequency<br>(#/10 min) | 4.3 | 2.86 | 38.4 | 6.08 | -5.07 | 8 | **0.001** |
| intruder agonistic frequency (#/10 min) | 7.7 | 2.71 | 32.3 | 3.58 | -5.50 | 8 | **0.001** |
| reproductive behavior latency (min) | 3.7 | 0.80 | 1.1 | 0.06 | 3.44 | 8 | **0.009** |
| resident agonistic latency (min) | 6.9 | 1.07 | 1.0 | 0.00 | 10.36 | 8 | **<0.001** |
| intruder agonistic latency (min) | 7.6 | 0.81 | 1.6 | 0.24 | 7.13 | 8 | **<0.001** |
| reproductive behavior intensity (0-3 scale) | 0.9 | 0.07 | 1.5 | 0.17 | 1.50 | 10 | **0.016** |
| resident agonistic intensity (0-3 scale) | 0.7 | 0.35 | 2.3 | 0.29 | 1.50 | 10 | **0.016** |
| intruder agonistic intensity (0-3 scale) | 0.5 | 0.22 | 2.0 | 0.13 | 0.00 | 10 | **0.008** |
| **Physical Variables:** |  |  |  |  |  |  |  |
| snout-vent length (mm) | 6.0 | 0.07 | 6.0 | 0.10 | 0.00 | 8 | 1.00 |
| body-minus-head mass (g) | 3.3 | 0.07 | 3.4 | 0.08 | -0.75 | 8 | 0.48 |
| testes mass (g) | 0.1 | 0.00 | 0.1 | 0.01 | -0.26 | 8 | 0.80 |
| testosterone (ng/ml) | 2.2 | 1.08 | 1.5 | 0.14 | 0.30 | 8 | 0.77 |
| estradiol (ng/ml) | 9.3 | 5.68 | 5.2 | 1.78 | 0.68 | 8 | 0.74 |
| progesterone (ng/ml) | 0.6 | 0.06 | 0.8 | 0.21 | -0.97 | 8 | 0.14 |
| glucocorticoids (ng/ml) | 7.0 | 3.14 | 4.8 | 0.59 | 0.25 | 8 | 0.81 |

### Boldness is associated with gene upregulation in the ventromedial hypothalamus

We measured baseline gene expression in bold and shy individuals across six brain regions that contribute to social decision-making and are functionally conserved across vertebrates (Kabelik et al., 2018; Newman, 1999; O’Connell and Hofmann, 2012, 2011; Thompson et al., 2008; Walton et al., 2010) (**Figure 2A**; Supplemental Excel File). We used a principal component analysis to visualize overall gene expression differences across brain regions and groups. Principal component (PC) 1 explained 22.8% of the variance, PC2 16.3%, and PC3 8.8% (**Figure 2B**). Brain regions separated significantly in PC2 (F_6,45_=463.4, p<2×10^−16^), but not by group or their interaction. Bold and shy groups separated significantly in PC3 (F_1,45_=4.727, p=0.0372), but not by brain region or their interaction. The number of differentially expressed genes across brain regions were relatively few (average of 8), with the exception of the ventromedial hypothalamus, where 608 genes were differentially expressed (**Figure 2A**). Across brain regions, only integrin alpha-10 (*itga10*) was consistently downregulated in bold individuals compared to shy individuals (**Figure 2C**), with the exception of the dorsal ventricular ridge where expression of this gene was not detected. We also specifically examined genes that are associated with bold or shy phenotypes in other taxa (**Figure 2D**), and every gene had significant expression differences across brain regions (See Supplementary Excel File for statistics). Only two genes had significant group by brain region interactions, where estrogen receptor alpha (*erα*) was elevated in the ventromedial hypothalamus of bold lizards (p=0.039) and corticotropin-releasing hormone (*crh*) was elevated in the hindbrain of shy males (p=0.028).

**Figure 2.**
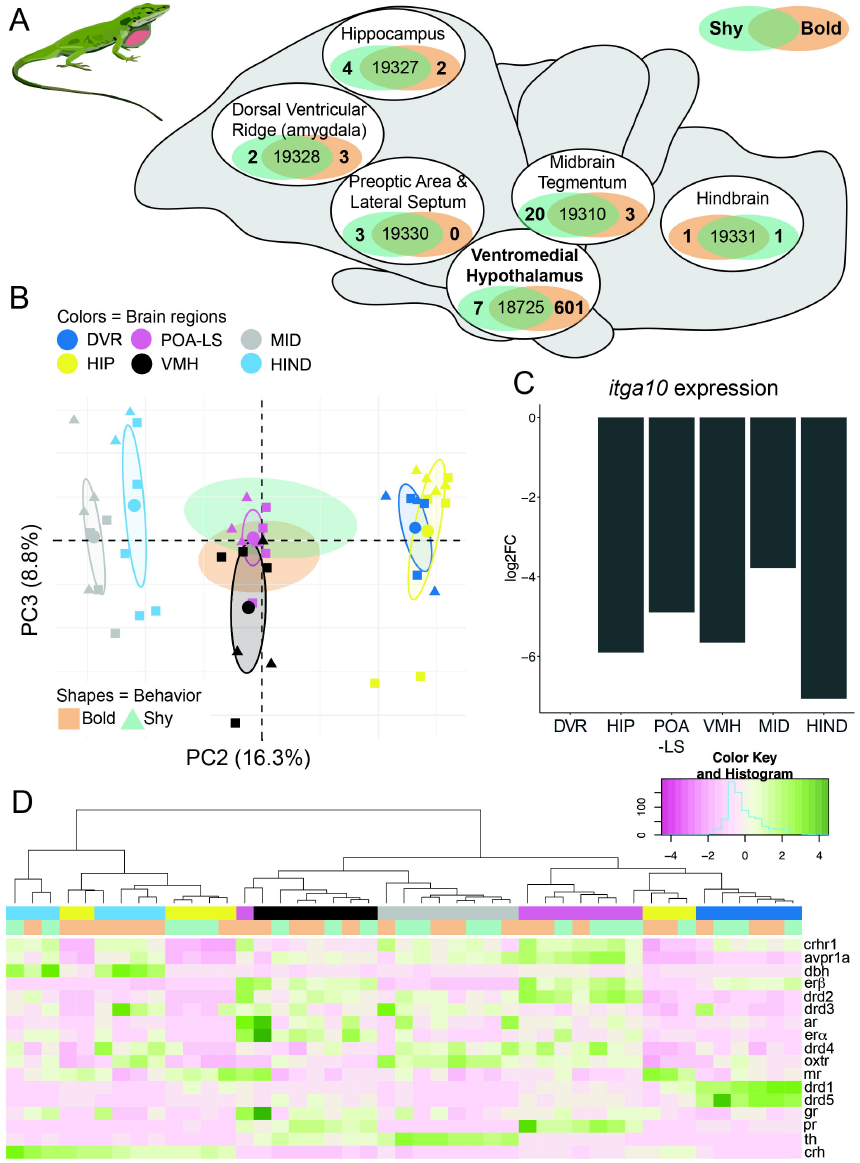
Baseline brain gene expression in bold and shy anoles. **(A)** RNA sequencing was used to quantify gene expression in six different brain regions in bold (orange) and shy (blue-green) individuals. The number of differentially expressed genes in each brain region is shown in the Venn diagrams. **(B)** A principal component analysis shows separation between brain regions (different colors) in PC2 and behavioral group (bold in squares and shy in triangles) in PC3. Ellipses show 95% confidence interval for each grouping. **(C)** Integrin alpha-10 (*itga10*) was downregulated in bold anoles across almost all brain regions. The log fold change (log2FC) is shown in bar plots. (D) A heatmap shows normalized expression of genes previously associated with bold and shy behaviors in other taxa (see text for references). Color bars below the dendrogram shows brain region and behavior groupings. The color key for the heat map shows green as increased expression and pink as decreased expression normalized within each gene.

Since the ventromedial hypothalamus had a drastically different pattern in baseline gene expression between bold and shy individuals, we explored these patterns in more detail (plots for all other brain regions are shown in Supplementary Figure 2). There were 601 genes upregulated and 7 downregulated in bold individuals compared to shy individuals (**Figure 3A**). While many differentially expressed genes are unannotated and labeled as “novel transcripts”, we noted several that have established roles in regulating behavior or have a log fold change of greater than 5 (**Figure 3B**). This includes the androgen receptor (*ar*, p=0.014) and the two subunits of the NMDA receptor, *grin1a* (p<0.001) and *grin2b* (p<0.001). Expression of integrin alpha-10 was downregulated in bold individuals (p=0.017), similar to other brain regions. Finally, some genes had a large fold change increase in bold individuals, including a potassium channel (*kcnh1-like*, p>0.001) and the secretin receptor (*sctr*, p=0.02), To further explore gene expression differences in an untargeted manner, we examined gene otology annotations for differentially expressed genes and found enriched molecular function of calcium channel activity (p=5.04×10^−4^). Indeed, at least seven voltage-gated calcium channel genes were upregulated in bold individuals (**Figure 3C**).

**Figure 3.**
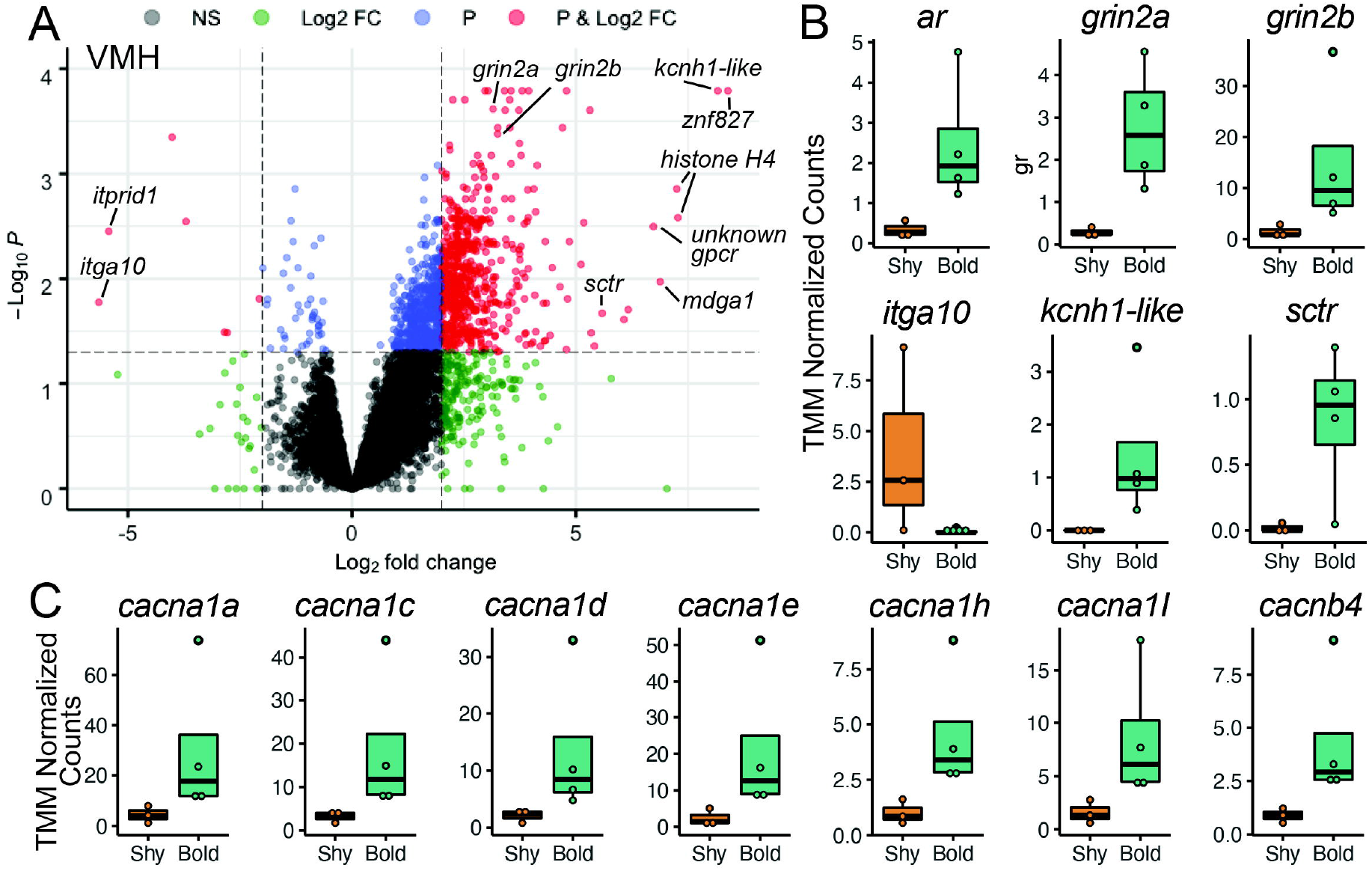
Baseline brain gene expression in bold and shy anoles highlights the ventromedial hypothalamus. **(A)** A volcano plot highlights the genes that are differentially expressed in the ventromedial hypothalamus, where the cutoff for significance (red dots beyond the dashed lines) is p<0.05 with false discovery rate correction and a log fold change of two or greater. **(B)** Genes previous linked to behavior and/or with large fold changes in expression and **(C)** voltage-gated calcium channels from the gene ontology analysis are shown as boxplots, with rectangles as the lower and upper quartiles (with the median as the line) and whiskers that indicate the maximum and minimum values; individual data points are shown as dots.

## Discussion

Most behaviorally linked gene expression studies examine changes resulting from participation in behavioral trials or the establishment of dominant or subordinate status within a social hierarchy. However, such comparisons make it difficult to ascertain what variables are predisposing animals to exhibit a bold or shy phenotype in the first place, as such studies will also detect gene transcription differences that result from the perception of conspecifics and performance of varied levels of behaviors toward other individuals. Hence, to remove perception-related and performance-based gene expression and focus on neural differences that predispose individuals toward bold or shy behavioral profiles, we examined male green anoles under baseline conditions. These anoles had been previously screened under three different social conditions and repeatedly within each condition. We found highly stable boldness phenotypes that transcended social context. That is, in relation to shy males, bold males tended to behave more quickly and to exhibit greater numbers and higher intensities of courtship behaviors within reproductive trials and of aggressive behaviors within agonistic trials. This correlation of social boldness across contexts is in line with expectations for consistent behavioral phenotypes or syndromes (Coppens et al., 2010; Koolhaas et al., 2010; Stamps and Groothuis, 2010). Thus, our gene expression analysis investigates neural correlates of general social boldness.

Because sequencing six brain regions from all 57 males in this study for the purposes of covariation analysis was beyond our means, we instead chose to focus on a comparison of individuals from the extreme ends of the normal boldness distribution found in this species. We reasoned that such a comparison would maximize our chances of detecting gene expression involved with the determination of social boldness. This approach has been used by other researchers (e.g., Thörnqvist et al., 2019) and is akin to the breeding of more bold or shy lines of rodents (Koolhaas et al., 2010) or zebrafish (Oswald et al., 2012; Wong et al., 2012a), although our study examines wild-caught animals whose phenotypic variance has been maintained solely though natural selection processes, rather than artificial selection. Future studies will examine specific genes of interest in specific brain tissues across larger sample sizes varying continuously in measures of social boldness, allowing for examination of covariance across a normal distribution.

### Evolutionary bases for stable boldness phenotypes

Behavioral boldness and stress coping styles appear to have strong heritable components (Ballew et al., 2017; Mont et al., 2018; Scherer et al., 2017) and variability of these traits within a population appears to be maintained by selective pressures (Koolhaas et al., 2010; Smith and Blumstein, 2010). Furthermore, as in this study, behavioral boldness in various species is stable, including across social contexts (hence eliciting the term behavioral syndrome) (Colléter and Brown, 2011; Koolhaas et al., 2010; Qu et al., 2018; Reaney and Backwell, 2007). This stability is likely due to underlying genetic correlations regulating behaviors across contexts (Oswald et al., 2013). In anole lizards, a finding of behavioral boldness stability across time was also recently reported in water anoles, *Anolis aquaticus* (Putman et al., 2019), supporting our results and a notion that behavioral boldness is a stable trait across anole species.

Studies of behavioral boldness have also shown that behavioral boldness is sometimes positively correlated with overall behavioral activity levels (Colléter and Brown, 2011; Smith and Blumstein, 2010; Wong et al., 2015). This association suggests the presence of shared behavioral circuitry beyond that for solely social behaviors. However, other studies find dissociations between locomotion and measures of boldness and stress responsiveness (Kanitz et al., 2019; Van Reenen et al., 2004), so such a correlation is not universal. General activity was not quantified in the present study, so future studies will need to determine whether bold male green anoles also exhibit greater general activity levels. Such a finding could provide evidence for increased behavioral display contributing to the increased frequency and decreased latency of displays during social interaction trials. However, a generally higher activity level in bold animals would less easily account for our finding of increased behavioral intensity in bold individuals, as activity levels in and of themselves would not determine whether an individual would, for instance, rather bite than flee from a conspecific during an agonistic trial. Thus, we suspect that although general behavioral activity may correlate with behavioral boldness in green anoles, social behavior regulation seems to be regulated, at least in part, by separate mechanisms.

### Social boldness phenotype is unrelated to body size and circulating steroid hormone levels

Some previous studies have found associations between behavioral boldness and measures of physical condition. These studies have focused on the enhanced physical abilities of bold individuals allowing them to better escape predators or obtain resources, thus providing fitness benefits (Mayer et al., 2016; Smith and Blumstein, 2010). A similar argument can be made that individuals in better physical condition may also be better combatants in agonistic encounters and thus bolder in such situations. However, while there were many behavioral differences between bold and shy individuals in this study, we did not find any group differences in body size. Furthermore, although body size has been found to correlate with boldness in a number of species (Adriaenssens and Johnsson, 2011; Brown and Braithwaite, 2004; Mayer et al., 2016), the direction of this correlation is not always consistent due to the underlying basis for the relationship. For example, smaller individuals of some species may need to leave shelter earlier to forage, while in other species a larger body size can allow individuals to avoid predation and more easily consume prey and thus allow larger individuals to more readily leave shelter. Additionally, other studies find no correlation between boldness and body size (Reaney and Backwell, 2007), presumably because no third variable drives such a relationship. The lack of relationship between body size and boldness in our study therefore suggests that boldness in male green anoles has fitness consequences largely independent of body size.

We initially hypothesized that there would be group differences in glucocorticoid hormone levels because a sub-section of studies that address behavioral boldness focus primarily on active versus passive coping styles to stressors, and some of these studies have found glucocorticoid differences between coping styles (Koolhaas et al., 2010; Sluyter et al., 1996). However, we found no relationship between circulating glucocorticoid levels and behavioral boldness in our study of male green anoles. Similarly, bold and shy male zebrafish also do not differ in baseline cortisol levels (Oswald et al., 2012), nor do rainbow trout (Gesto, 2019), sea bass (Alfonso et al., 2019), or pigs differing in coping style (Kanitz et al., 2019). The findings in rodents are mixed, with some studies finding correlations between basal glucocorticoid levels and measures of boldness, while others do not (Koolhaas et al., 2010). However, we must also consider additional variables such as time and context of hormone sampling. Our hormone measures were from diurnal plasma samples, and thus we cannot exclude the possibility of group differences in basal glucocorticoid release at other points in the circadian cycle, such as those seen in zebrafish (Tudorache et al., 2018) and mice (Veenema et al., 2004, 2003). Our hormone measures were also from baseline plasma samples, and thus we cannot exclude the possibility of group differences in stress-induced glucocorticoid release. This latter scenario may be possible given that we detected elevated *crh* in the hindbrain of shy males, although we found no such difference in hypothalamic regions. Much variation exists in the literature, with various findings across species, and even within single species, suggesting that in some cases relationships exists between basal and/or stress-reactive glucocorticoid levels and behavioral boldness, while in other cases such relationships are absent (Koolhaas et al., 2010; Steimer et al., 1997). The concept of an (at least partial) dissociation between behavioral boldness and stress responsiveness in some species is supported by studies in rats, cattle, and pika (Qu et al., 2018; Steimer et al., 1997; Van Reenen et al., 2005, 2004). Research on stress-reactivity in green anoles is necessary to clarify whether such a dissociation is present in this species.

Organizational effects of steroid hormones on social behavior phenotype are present in some species (Koolhaas et al., 2010) and may also be present in green anoles. However, we find no evidence for activational effects of steroid hormones on determining boldness phenotype in anoles, despite their presumed involvements in other aspects of behavioral motivation and display. We had expected that testosterone, estradiol, or progesterone levels could correlate with group differences in behavioral boldness because of the known involvement of these hormones in the regulation of lizard social behaviors. In male tree lizards, circulating testosterone correlates with aggression (Kabelik et al., 2006) and testosterone and progesterone treatments causally promote aggression (Kabelik et al., 2008b; Weiss and Moore, 2004). Similarly, in male side-blotched lizards, circulating testosterone levels are higher in the more aggressive morph (Sinervo et al., 2000), and the anti-androgen cyproterone acetate has been found to reduce display behaviors in male brown anoles, toward both conspecific males and females (Tokarz, 1995). Studies examining other vertebrate taxa and focusing on behavioral boldness have found mixed associations with circulating steroid hormone levels. For example, testosterone treatment in African striped mice (*Rhabdomys pumilio*) increases boldness behavior (Raynaud and Schradin, 2014), and acute 17a-ethinylestradiol (an estrogen mimic) decreases boldness behavior in Siamese fighting fish (*Betta splendens*). However, aromatase inhibitors decrease boldness in female Siamese fighting fish, highlighting a role for local hormone synthesis within the brain. When performing our targeted gene expression analyses, we found that estrogen receptor alpha expression is increased in the ventromedial hypothalamus of bold males, suggestion estrogenic action in this brain region may be important in this context. Functional manipulations would be required to determine a causal role for steroid hormone action or synthesis in the brain and any relationships with individual variation in behavior. Another example of how steroid hormones may influence boldness is via androgen receptor expression. We further address this hypothesis below, in relation androgen receptor differences between groups.

### Differentially expressed genes and a regulatory role for the ventromedial hypothalamus

When comparing forebrain, midbrain, and hindbrain regions of bold versus shy male green anole lizards, we found the greatest number of differentially expressed genes within the ventromedial hypothalamus, a node within the social decision-making neural network (O’Connell and Hofmann, 2011). We were initially surprised to find so few genes differentially expressed within other brain regions, especially in regions such as the preoptic area, given its established role in regulating male anole sexual behavior (Crews and Morgentaler, 1979; Kabelik et al., 2013; Morgantaler and Crews, 1978; O’Bryant and Wade, 2002; Wheeler and Crews, 1978). However, the patterns here represent a constitutive state, rather than a response to a behavioral stimulus that would be expected to induce a greater change in gene expression across brain regions. The ventromedial hypothalamus has been implicated in the regulation of both reproductive and agonistic behaviors, making it a logical location for regulation of general social behavior boldness. For instance, both copulatory and agonistic conditions have been shown to upregulate markers of neural activity within the Syrian hamster ventromedial hypothalamus, although copulation tends to induce more c-Fos expression (an immediate early gene product and proxy marker of neural activity) in the medial portions of the nucleus, while agonistic situations tend to increase c-Fos in the lateral ventromedial hypothalamus (Kollack-Walker and Newman, 1995). Additionally, stimulation of the lateral portions of the rat ventromedial hypothalamus has been shown to elicit aggressive responses (Kruk, 1991). The ventromedial hypothalamus therefore seems a likely social behavior integration center, owing to its regulatory role in various types of social behavior expression.

In the ventromedial hypothalamus, we observed increased gene expression of a few neuromodulators with strong ties to behavior. For example, although we did not find significant differences in testosterone levels between bold and shy individuals, we did find an increase in androgen receptor expression in bold individuals. This finding is exciting given that androgen receptor presence within the ventromedial hypothalamus occurs within its dorsolateral aspect (Rosen et al., 2002), the same region of the ventromedial hypothalamus that has been previously linked to the expression of aggressive behavior in male tree lizards (Kabelik et al., 2008a). Although they did not examine the ventromedial hypothalamus, Hattori and Wilczynski (2014) found increased androgen receptor expression in the preoptic area and anterior hypothalamus of green anoles that won agonistic encounters and became dominant to a same-sex conspecific, further suggesting a role of the androgen receptor in mediating social interactions. Further studies will be necessary to determine whether social interactions likewise mediate androgen receptor expression within ventromedial hypothalamus.

We also found an increase in expression of the secretin receptor in bold individuals. Secretin is primarily known for its role in regulating neurodevelopment and memory function via effects on synaptic plasticity (Wang and Zhang, 2020). Synaptic plasticity, learning, and memory formation have been linked to stress coping styles and behavioral boldness (Bolhuis et al., 2004; Coppens et al., 2010; Delclos et al., 2020; Wong et al., 2015). Furthermore, secretin has also been shown to regulate anxiety and associated behaviors (Nishijima et al., 2006; Wang et al., 2019), as well as to modulate activity of reproductive circuitry (Csillag et al., 2019). More specifically, secretin regulates GABAergic transmission to gonadotropin-releasing hormone-producing cells (Csillag et al., 2019), as well as affecting the firing rate of over 50% of examined paraventricular hypothalamus neurons in rat *in vivo* studies (Chen et al., 2013). These studies suggest a role for secretin as a widespread modulator of neural function, and as such, secretin may also regulate social behavior via modulation of neuronal activity within the ventromedial hypothalamus.

A prominent category of differentially expressed genes within the ventromedial hypothalamus, as revealed by the gene otology analysis, was voltage-gated calcium channels. Voltage-gated calcium channels regulate both intracellular calcium levels as well as general neuronal excitability, and have been linked to a number of neuropsychiatric symptoms in humans, including bipolar disorder, depression, and attention deficit hyperactivity disorder (Kabir et al., 2017). In addition to voltage-gated calcium channels, we also found that expression of ligand-gated calcium channel NMDA subunits is upregulated in bold individuals. NMDA receptor regulation has been linked to behavioral boldness in birds (Audet et al., 2018), suggesting the tuning of calcium channel expression may be a conserved feature of behavioral variability. Indeed, increased intracellular calcium levels induce signaling cascades that can lead to changes in transcription, such as the phosphorylation of cyclic adenosine monophosphate response element binding protein (CREB), which has been linked to synaptic, neuronal, and behavioral plasticity (Hofmann, 2003). In tree lizards, the dorsolateral portions of the ventromedial hypothalamus show increased neural activity as measured by an increase in pCREB following an agonistic encounter (Kabelik et al., 2008a). Thus, a number of neural activity-regulating channels differ in baseline gene expression between bold and shy males, highlighting neuronal excitability in the ventromedial hypothalamus as a contributor to stable individual variation in behavior.

Across most brain regions, integrin alpha-10 was the one gene consistently downregulated in bold individuals. Integrins are typically associated with neuronal development, as they can detect and transmit mechanical force on extracellular matrices into an intracellular signal (Takada et al., 2007). The role of integrin alpha-10 is not well understood, especially in the context of behavior. Integrin alpha-10 in humans is associated with the 1q21.1 chromosomal region, which when deleted leads to thrombocytopenia absent radius (TAR) (Brunetti-Pierri et al., 2008). Genome wide association studies in humans has also linked integrin alpha-10 to bipolar disorder (Pedroso et al., 2012). Thus, a role for integrin alpha-10 in natural variation in behavioral strategies beyond that of human disease is a promising avenue of future research.

Other neuromodulators previous linked to coping styles in fish and mammals were not differentially expressed between bold and shy male green anoles, including dopamine, nonapeptides, and the glucocorticoids. We did find increased corticotropin releasing hormone expression in the hindbrain of bold males, but this should be interpreted with caution due to our low sample size. Bold zebrafish tend to have increased D2 receptor expression (Thörnqvist et al., 2019) and shy sea bass have increased glucocorticoid-related gene expression (Alfonso et al., 2019), but these studies analyzed whole brain gene expression patterns rather than specific brain regions. It is possible that D2 receptor or glucocorticoid-related gene expression may also differ between bold and shy male green anoles within brain regions not examined in our study. Studies on artificially selected lines of bold and shy zebrafish have found that shy lines have elevated expression of glucocorticoid-related genes in pooled hypothalamus/midbrain/optic tectum samples (Oswald et al., 2012) and neurometabolism genes in whole brain tissue (Wong et al., 2015). We did not find strong evidence of group differences in expression of these genes, which could be due to species differences in neural mechanisms or differences in design as the fish study used artificially selected lines. Glucocorticoid receptors as well as nonapeptide receptors have also been associated with coping styles in mammals, where Kanitz et al. (2019) found differential brain gene expression in piglets exposed or not exposed to a stressor, although only glucocorticoid receptor expression differences in the hypothalamus were apparent between groups. In this same piglet study, V1a receptor differences were only present as interaction effects with stress treatment and this was only observed within the hypothalamus. The lack of differential expression of these genes in hippocampal and amygdalar tissues is consistent with our findings, while the findings in the hypothalamus are more difficult to interpret since the entire hypothalamus was sampled in the pig study. Overall, the general patterns of gene expression across taxa are difficult to compare given differences in behavioral paradigms and tissue sampling. More consistent behavioral paradigms and brain region sampling across taxa are needed for more fair comparisons.

## Conclusion

Our study focuses on constitutive differences across bold or shy individuals by isolating baseline hormone levels and brain gene expression profiles that influence social boldness predisposition, rather than those reflecting the results of social interaction and behavioral execution. We found that correlated behavioral traits were not driven by differences in body size or steroid hormone levels, as these were consistent across treatment groups. Instead, brain gene expression differences strongly relate to social boldness and likely reflect variables involved in the neural circuitry that regulates social boldness. Specifically, we found that baseline differences between bold and shy males were associated with gene expression in the ventromedial hypothalamus, where expression of calcium channels, the androgen receptor, and the secretin receptor were increased in bold individuals. We suggest that studies should include examination of the ventromedial hypothalamus as a potential regulator of social behavior boldness in reptiles as well as across other vertebrate taxa.

## Acknowledgements

DR and LAO acknowledge that Stanford University resides on the ancestral and unceded land of the Muwekma Ohlone Tribe.

## Data Accessibility

Data from the behavior and hormone analyses, as well as RNA sequencing analysis, including count matrices, GO enrichment analyses, and differential expression statistics, are available in the Supplementary Excel File. Raw sequencing reads are available on the Sequence Read Archive (submission pending acceptance).

